# Cis-regulatory chromatin contacts form de novo in the absence of loop extrusion

**DOI:** 10.1101/2025.01.12.632634

**Authors:** Nicholas G. Aboreden, Han Zhao, Fengnian Shan, Fuhai Liu, Haoyue Zhang, Gerd A. Blobel

## Abstract

NIPBL promotes chromatin loop extrusion by the cohesin complex until it stalls at convergently oriented CTCF sites, leading to the formation of structural loops. However, to what extent loop extrusion contributes to the establishment vs maintenance of *cis*-regulatory element (CRE) connectivity is poorly understood. Here, we explored the de novo establishment of chromatin folding patterns at the mitosis-to-G1-phase transition upon acute NIPBL loss. NIPBL depletion primarily impaired the formation of cohesin-mediated structural loops with NIPBL dependence being proportional to loop length. In contrast, the majority of CRE loops were established independently of loop extrusion regardless of length. However, NIPBL depletion slowed the re-formation of CRE loops with weak enhancers. Transcription of genes at NIPBL-independent loop anchors was activated normally in the absence of NIPBL. In sum, establishment of most regulatory contacts and gene transcription following mitotic exit is independent of loop extrusion.

## INTRODUCTION

Mammalian genomes are organized in a nonrandom, multi-layered manner within the nucleus. Spatial organization is intimately connected to gene regulation as, for example, distal transcriptional enhancers are in physical proximity with their promoters^1,2^. An important player in the establishment and maintenance of genome structure is the cohesin complex, which actively extrudes the chromatid until it encounters convergently oriented CTCF sites, leading to the formation of looped contacts, also referred to as structural loops^3–6^. These loops give rise to topologically associating domains (TADs), which promote regulatory interactions within their boundaries and disfavor contacts across them^7–11^.

TADs and structural loops are dynamic entities, persisting for mere minutes and are thus subject to constant turnover^12,13^. Cohesin activity is modulated by factors such as WAPL/PDS5^14–18^, which facilitate the release of cohesin from chromatin, and NIPBL, a cohesin activator. NIPBL was identified in Drosophila as a facilitator of long-range enhancer-promoter communication^19^ and is required for cohesin-mediated loop extrusion^14,20^. While NIPBL has been thought of as a cohesin loader onto chromatin ^21–25^, recent evidence suggests an additional role, in which NIPBL primarily enhances the processivity of cohesin extrusion^15,26–30^. In-vitro biochemical experiments demonstrated that cohesin’s ATPase activity is greatly stimulated by NIPBL, providing a mechanistic explanation for how NIPBL enhances cohesin processivity^27,28^.

Despite the established roles of CTCF and cohesin in shaping genome architecture, the contribution of the CTCF/cohesin machinery to cis-regulatory element (CRE) connectivity is limited. The transcription of most genes and the interaction between numerous CREs persist following acute CTCF or cohesin depletion^18,31–36^, pointing at the existence of CTCF/cohesin-independent mechanism that connect CREs. However, to what extent cohesin-mediated loop extrusion is necessary for the initial establishment of CRE interactions versus their maintenance, remains to be explored more deeply.

Mitosis is characterized by a dramatic reorganization of chromatin architecture, with most measurable features such as compartments, TADs, and chromatin loops being transiently disrupted, only to be rapidly re-established during G1 entry^37–41^. This unique transition period offers a powerful opportunity to study the establishment of architectural features and the factors involved.

The study of cohesin in chromatin architecture during mitotic exit is complicated by its essential role in maintaining sister chromatid cohesion in pro-metaphase. Therefore, to explore the role of loop extrusion in establishing chromatin architecture without disrupting mitosis we depleted NIPBL. Specifically, we performed Hi-C in cells that were depleted of NIPBL during mitosis, followed by synchronous release into G1. We found that most, but not all, cohesin-dependent structural loops failed to form. The dependence of structural loops on NIPBL was associated with loop length - longer loops were more sensitive to NIPBL depletion than shorter ones. Importantly, most CRE loops and gene transcription were re-established normally. Hence, loop extrusion only plays a minor role in establishing CRE connectivity implying alternative mechanisms, such as transcription (co-)factor clustering by which CREs are formed in the 3D space.

## RESULTS

### NIPBL supports the formation of TADs and stripes

To test the requirement of NIPBL for the de novo formation of chromatin architecture genome-wide, we tagged endogenous NIPBL bi-allelically with an 5-Ph-IAA-inducible degron^42,43^ and a HALO tag in the G1E-ER4^44^ erythroblast cell line using CRISPR-mediated gene editing. We introduced an osTIR2-expressing construct into the cells to allow for 5-Ph-IAA-inducible NIPBL degradation. NIPBL was virtually completely degraded upon 4 hrs of 5-Ph-IAA treatment, as measured by western blot in cell lysates (Fig. 1a). To investigate the impact of NIPBL loss specifically during the mitosis-to-G1 transition, a period when chromatin architecture is rebuilt, we treated NIPBL-AID cells with 5-Ph-IAA for 4 hours during nocodazole-induced prometaphase arrest and continued the 5-Ph-IAA treatment upon nocodazole release into G1. To analyze the temporal dynamics of chromatin architecture formation upon NIPBL depletion, we collected 3 timepoints during G1 entry; two in early G1 (45min and 1hr) and one in late G1 (4hr) (Fig. 1b). 2N cells were purified based on DNA content via DAPI staining to ensure appropriate release into G1, followed by in situ Hi-C.

**Fig. 1:**
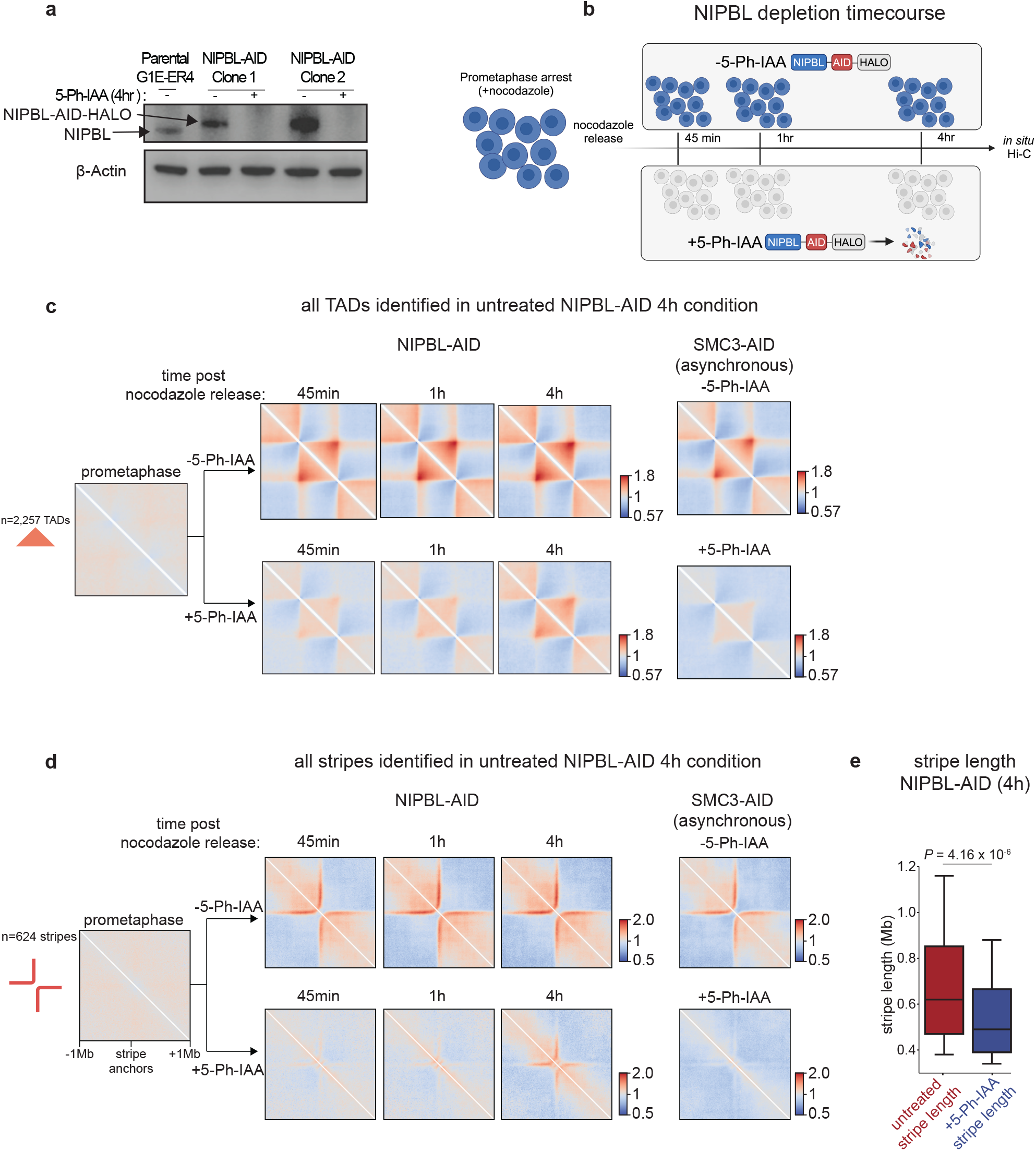
NIPBL is required for domain and stripe formation after mitosis. **a** Western Blot in asynchronous NIPBL-AID G1E-ER4 clones upon 4hr of 5-Ph-IAA treatment. **b** Schematic of NIPBL depletion time course during G1 entry. **c** Pile-up plots showing average signals for TADs identified in untreated 4h release condition. **d** Pile-up plots showing average signals for stripes identified in untreated 4h release condition. **e**. Stripe length for stripes identified in untreated 4h release condition compared to those identified in 5-Ph-IAA-treated 4h release condition.

We began testing the role of NIPBL in establishing chromatin architecture by analyzing the re-formation of TADs. Using the hicFindTads function from HiCExplorer^45^, we identified a total of 2,257 TADs in control cells in late G1 (4 hours). NIPBL loss globally impaired TAD re-formation (Fig.1c). However, residual TADs exhibited a progressive strengthening as cells advanced through G1, possibly due to NIPBL-independent extrusion by cohesin. To validate that the NIPBL-dependent TADs are indeed cohesin-dependent structures, we analyzed Hi-C data from asynchronously growing (mostly interphase) G1E-ER4 cells bearing AID tagged SMC3 (cohesin) alleles, treated with 5-Ph-IAA for 4 hours^46^. SMC3 depletion led to robust disruption of these TADs, confirming that the domains in question are indeed cohesin-mediated.

To independently assess NIPBL’s influence on cohesin-mediated loop extrusion, we analyzed the formation of “stripes” or “flares” in NIPBL-depleted cells using the Stripenn algorithm^47^. Stripes, which appear as horizontal or vertical lines on Hi-C maps, represent loop extrusion intermediates^6,36,48^. We found that stripes were virtually absent at early G1 timepoints following mitotic NIPBL depletion and began to appear by late G1, although they remained less intense and shorter than those observed in control cells (Fig. 1d,e). Control cells exhibited robust stripe formation in early G1, reflecting strong cohesin activity upon immediate entry into G1. Thus, mitotic NIPBL depletion impairs cohesin activity, potentially by reducing its extrusion rate and/or efficiency.

### Mitotic NIPBL loss strengthens compartmentalization

Prior reports suggest that cohesin-mediated loop extrusion counteracts genome compartmentalization^20,49,50^. To test whether NIPBL loss enhances the segregation of A and B-type compartments during mitotic exit, we measured genome-wide compartmentalization as cells entered G1 in the presence or absence of NIPBL. NIPBL loss strengthened compartmentalization as cells progressed into G1. This was evidenced by a sharper distinction between A and B compartments and increased intra-compartment versus inter-compartment interactions (Extended Data Fig. 1c-e).

### CRE loops can form independently of loop extrusion

CRE loops can persist upon acute depletion of CTCF or cohesin in interphase, suggesting that alternative mechanisms maintain CRE connectivity^34,51–53^. However, whether cohesin-mediated extrusion is required for the initial establishment of CRE loops upon mitotic exit remains to be directly tested. Indeed, it has been proposed that cohesin may be more essential for the establishment of de novo loops/gene expression profiles during cell state transitions than it is for maintaining them^54^. We identified loops using the cooltools.dots function in untreated 4hr G1 and 5-Ph-IAA-treated 4hr G1 samples. We quantified loop strength for each loop in each condition by measuring the observed contact frequency within the center (peak) pixel divided by a locally adjusted expected value. We then assigned a log2FC value comparing NIPBL replete and depleted conditions for each loop and used a log2FC cutoff of −/+ 0.5 to define differential loops.

We classified loops into three categories based on our prior annotations^41^: 1) structural loops – CTCF/RAD21(cohesin) peaks at both anchors and CREs present at one or no anchor, 2) CRE loops – CREs at both anchors and CTCF/RAD21 peaks at one or no anchor, 3) dual function loops – CREs at both anchors and CTCF/RAD21 peaks at both anchors (Fig. 2a). “Anchors” we defined as 20kb windows around the centers of the contacts. NIPBL depletion predominantly impaired the formation of structural and dual-function loops. Specifically, at late G1, 56% of structural loops failed to achieve maximal intensity while most of the rest were delayed in their formation (Fig. 2b-d). Similar results were observed for dual function loops. In stark contrast, 76% of CRE loops formed at similar rates and achieved normal intensities in the absence of NIPBL.

**Fig. 2:**
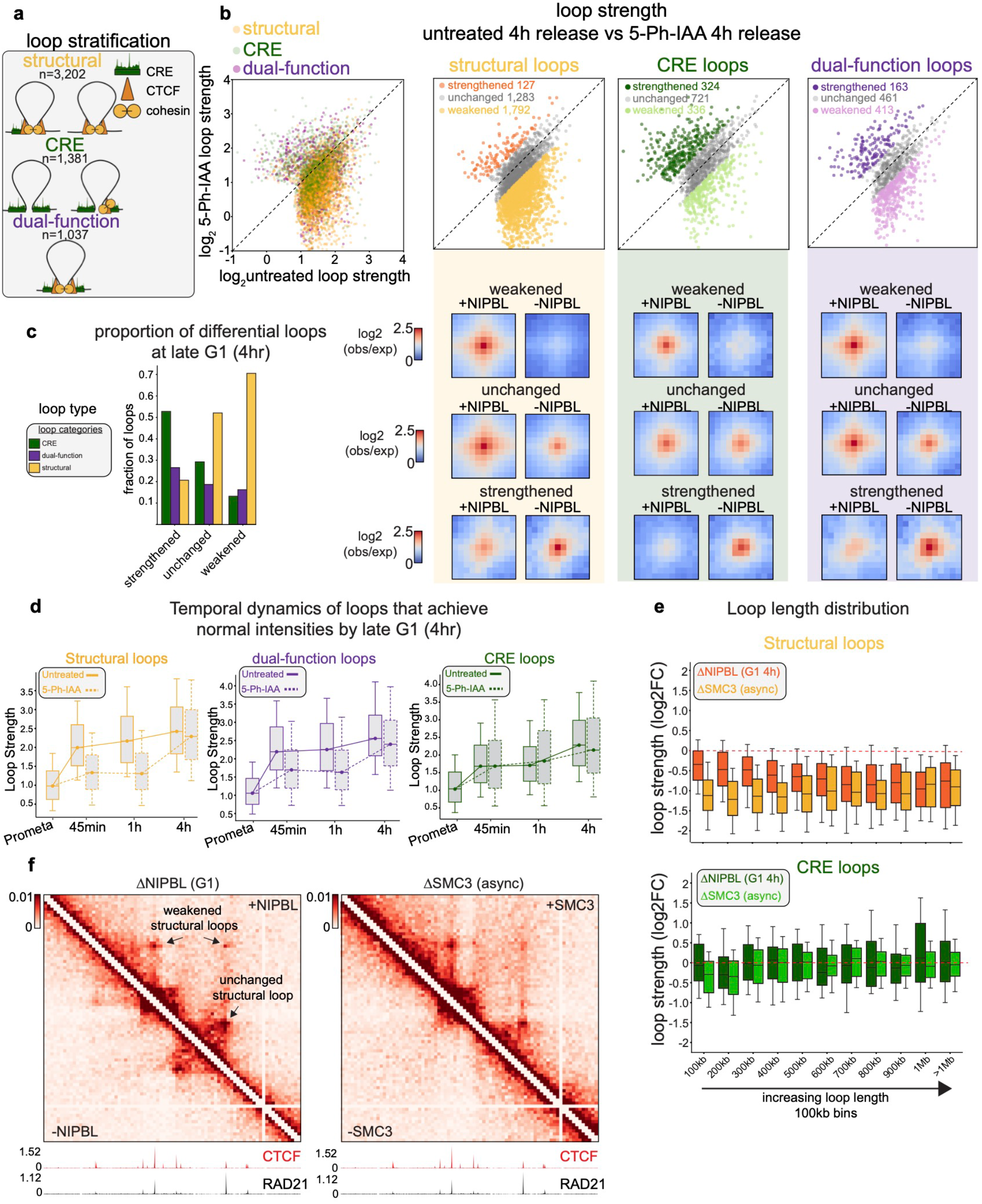
Chromatin loops display distinct responses to NIPBL loss. **a**. Schematic of loop stratification strategy. **b**. Differential loop analysis between untreated and 5-Ph-IAA-treated 4h release conditions. (Top) scatter plots showing individual loop responses to NIPBL depletion. (Bottom) APA plots showing average signals for weakened, unchanged, and strengthened loops. **c**. Fraction of strengthened, unchanged, and weakened loops that are classified as CRE, structural or dual-function. **d**. Temporal dynamics of loop strength for loops that achieve normal intensities by late G1 upon NIPBL depletion. **e**. Change in loop strength due to either NIPBL depletion (4h nocodazole release) or SMC3 depletion (asynchronous) for structural and CRE loops. Loops stratified by their length. **f**. Hi-C contact matrices of representative region containing structural loops with variable reliance upon NIPBL. (Left) NIPBL-AID at 4h release condition. (Right) SMC3-AID in asynchronous cells before/after 4h 5-Ph-IAA treatment.

If an extrusion process is involved in long range contact formation, loop length and dependence on NIPBL are expected to correlate. Indeed, NIPBL’s influence on cohesin-dependent structural loops was proportional to loop length (longer structural loops tended to be more sensitive to NIPBL depletion than shorter ones) (Fig. 2e,f). Interestingly, this trend did not exist for CRE loops which could be robust to NIPBL loss across length scales (Fig. 2e). Thus, CRE loops can be established independently of NIPBL/cohesin-mediated loop extrusion.

### Normal activation of most genes in G1 in the absence of NIPBL

To test whether loop extrusion is required to establish transcription, we measured nascent transcription by TT-seq in cells depleted of NIPBL during mitosis and released into G1 phase for either 45 minutes or 4 hours. NIPBL loss resulted in relatively modest gene expression changes, with 187 genes differentially expressed at the 45-minute timepoint and 188 at 4 hours (log2FC cutoff 1/−1, Padj < 0.05) (Fig. 3a). 93 differentially expressed genes (DEGs) were shared between both timepoints (Fig. 3b). Genes known to be activated in early G1 such as Fos and Jun were not differentially expressed at either timepoint, suggesting that NIPBL depletion does not interfere with mitotic release and/or G1 entry.

**Fig. 3:**
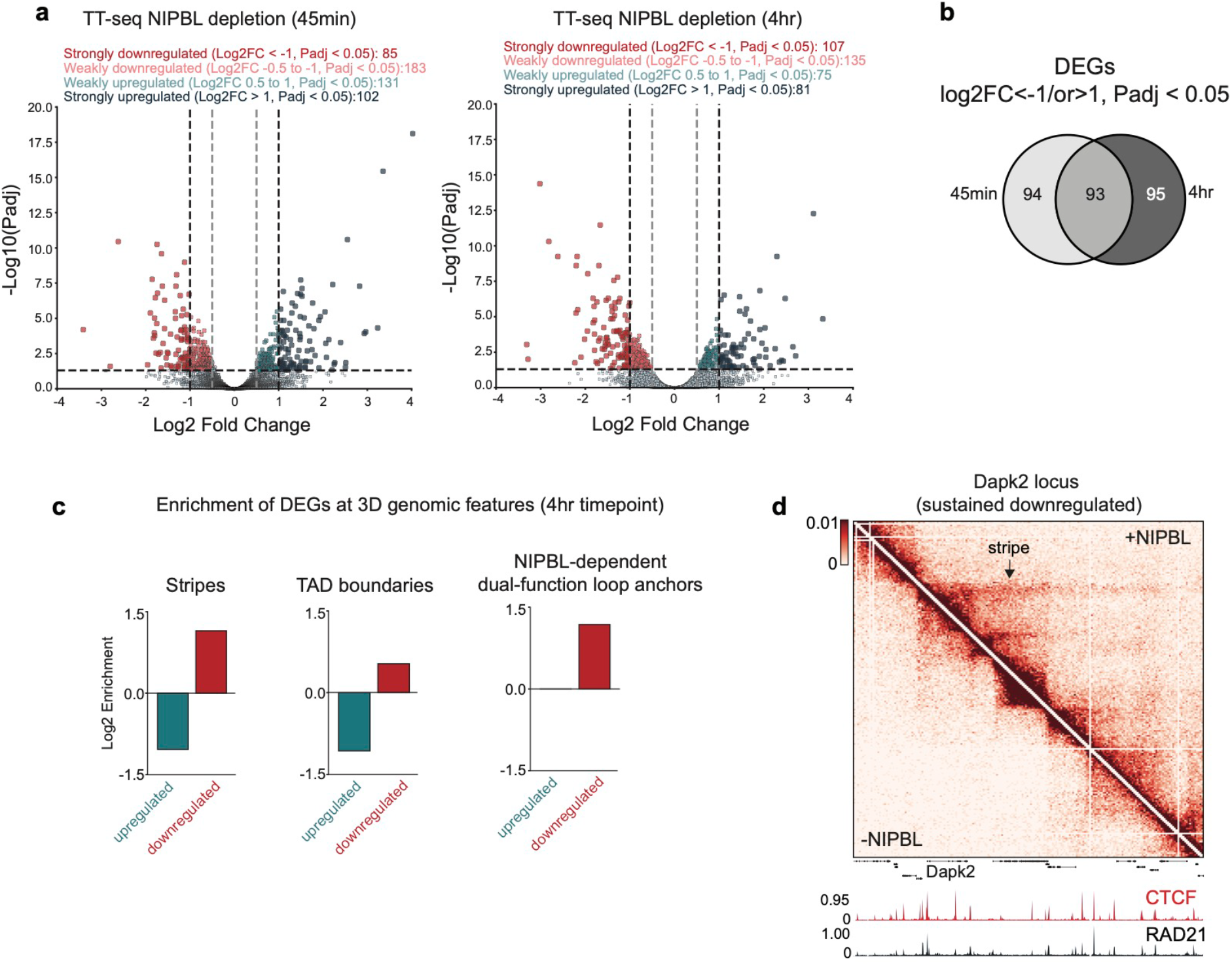
Modest transcriptional changes upon NIPBL loss. **a**. Volcano plots showing differential gene expression upon mitotic NIPBL depletion and release into G1 for 45min (left) and 4h (right). **b**. Overlap of differentially expressed genes between 45min and 4h timepoints. **c**. Enrichment of differently expressed genes (4h timepoint) at stripe anchors, TAD boundaries and NIPBL-dependent dual-function loop anchors. **d**. Hi-C contact matrices in the presence and absence of NIPBL (4h G1 release) of representative region containing NIPBL-dependent stripe anchored on NIPBL-dependent gene.

Of note, NIPBL-dependent genes did not overlap with NIPBL-dependent CRE loop anchors (not shown). In contrast, these genes were enriched at architectural stripes, TAD boundaries, and NIPBL-dependent dual-function loop anchors (Fig. 3c-d). Thus, the small subset of NIPBL-dependent genes likely relies on structural loops (including dual function loops) to promote CRE proximity, explaining their dependence on NIBPL.

One caveat to this analysis is that the genome undergoes a global spike in transcriptional activity during the M-G1 phase transition^55^. This spike may mask NIPBL-dependent changes in gene expression. Nevertheless, our data suggest that transcription is re-established mostly normally, despite loop extrusion inhibition.

### LDB1- and YY1-dependent loops can form in the absence of NIPBL

What mechanisms drive CRE connectivity in the absence of NIPBL? To this end, we focused on LDB1 and YY1-two architectural factors previously shown to have cohesin-independent roles^52,53,56^. Using Micro-C datasets generated in asynchronous LDB1-AID and YY1-AID G1E-ER4 cells, we tested whether loops dependent on LDB1 or YY1 were affected by mitotic NIPBL depletion. Focusing on loops with CREs in both anchors (this includes dual-function and CRE loop categories), we found that the majority of loops dependent on either LDB1 or YY1 were able to form in the absence of NIPBL (Fig. 4a,b and Extended Data Fig. 2a,b). However, a subset of LDB1/YY1 loops were NIPBL-dependent and tended to be closely flanked by encompassing structural loops (consistent with previous reports of cohesin-dependent CRE loops^51,53^ Fig. 4c and Extended Data Fig. 2c). Additionally, another subset of LDB1/YY1 loops exhibited delayed re-formation upon NIPBL depletion, but eventually recovered in late G1. Thus, NIPBL is dispensable for the establishment of the majority of LDB1/YY1-mediated regulatory interactions but may facilitate the timely formation of a subset of such loops.

**Fig. 4:**
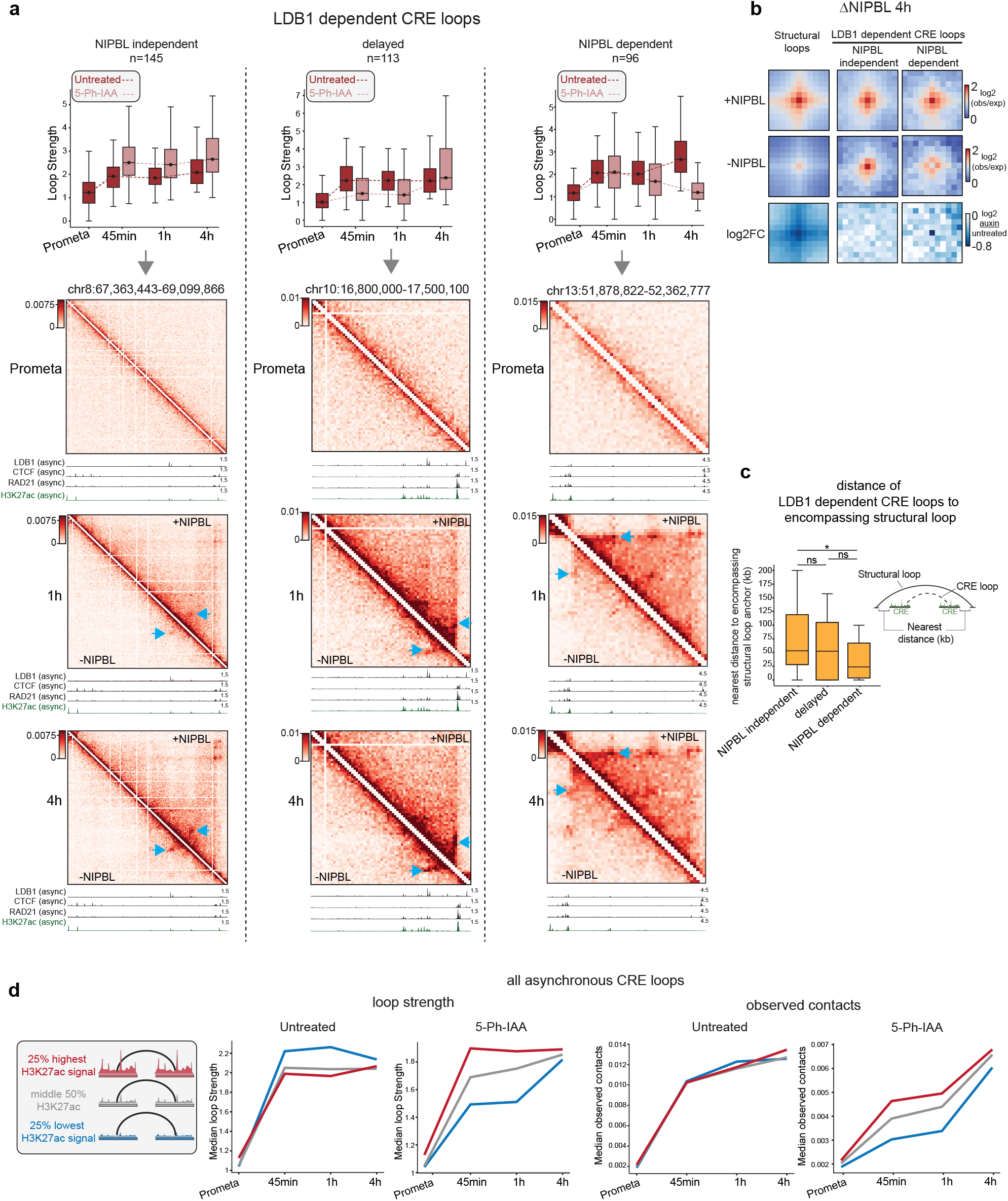
LDB1-dependent loops can form after mitosis in the absence of NIPBL. **a**. (Top) Loop strength values during G1 entry for LDB1-dependent CRE loops classified as NIPBL independent, delayed, and NIPBL dependent. (Bottom) Hi-C contact matrices of representative regions containing LDB1-dependent loops that are independent, delayed, and dependent on NIPBL. **b**. APA plots showing average signal for LDB1-dependent CRE loops in the presence and absence of NIPBL (4h G1 condition). **c**. Distance of LDB1-dependent CRE loops to encompassing structural loops. Only loops with identified structural loops encompassing them are shown. **d**. Loop strength (right) and observed contacts (left) for all asynchronous CRE loops during G1 entry in the presence and absence of NIPBL. Loops are stratified by the average H3K27ac signal at their anchors.

An important technical consideration is that loop strength was calculated by dividing the observed contacts within the loop’s peak pixel by a locally adjusted expected value. Since NIPBL depletion disrupts most TADs, global background contact frequencies are lower (Extended Data Fig. 2d). Regardless, the strength (observed over locally-adjusted expected signal) of LDB1/YY1-dependent/NIPBL-independent loops remains unchanged or even increases. Measured increases upon NIPBL depletion may be genuine or simply result from the peak pixels appearing stronger over the reduced local background.

To further test whether YY1 and LDB1 mediated interactions can form in the absence of NIPBL, we used an independent approach by generating pairs of YY1, LDB1, or RAD21 (as a control) ChIP-seq peaks. We created all possible pairs of peaks within a maximum distance of 500kb from each other and averaged the contact frequency between these pairs in the presence and absence of NIPBL (4hr post nocodazole release). As expected, contacts between RAD21 sites were weakened by NIPBL depletion. Contacts between LDB1 and YY1 peaks were retained to a greater extent, even though background contact frequencies surrounding the “loop” center were reduced (Extended Data Fig. 2e). This indicates that LDB1 and YY1-driven interactions can be established independently of NIPBL/cohesin, providing an alternative mechanism of loop formation.

### NIPBL Facilitates Rapid Formation of Loops Involving Weak CREs During Mitotic Exit

CRE proximity to structural loops can render CRE loops dependent on NIPBL (Fig. 4c and Extended Data Fig. 2c). However, prior work also suggested that CRE loops involving weak enhancers tend to rely more on support by structural loops than those involving stronger enhancers^51^. To investigate whether CRE activity is associated with loop dependence on NIPBL, we grouped all asynchronous CRE loops into three categories based on H3K27ac signal intensity at their anchors: top 25%, middle 50%, and bottom 25%. Loops with the lowest H3K27ac levels were most affected by NIPBL depletion and were delayed in their formation, while loops with the highest H3K27ac levels remained largely unaffected and formed quickly during mitotic exit (Fig. 4d).

Under NIPBL replete conditions, CRE loops of all three groups were established at comparable rates. Thus, weak enhancers tend to rely on extrusion to quickly establish loops during mitotic exit even though many of these still eventually form in late G1.

## DISCUSSION

The mitosis-to-G1 phase transition offers a unique opportunity to study the mechanisms driving chromatin architecture formation. However, disrupting cohesin during the mitosis-to-G1 phase transition is complicated due to its essential role in sister chromatid cohesion and segregation. This challenge has precluded direct assessment of the role of cohesin-mediated loop extrusion in re-establishing chromatin architecture. To circumvent this challenge, we used mitotic degradation of NIPBL, a key activator of cohesin. This approach allowed us to selectively impair cohesin’s extrusion activity during mitotic exit, providing valuable insights into NIPBL’s role in regulating cohesin and the necessity of loop extrusion for re-establishing chromatin architecture.

Long range structural loops are dependent on NIPBL while shorter ones can form but with delayed kinetics. The relationships of loop size and kinetics are consistent with a model in which NIPBL enhances cohesin’s extrusion processivity, allowing it to extrude further and faster. This is consistent with in vitro experiments demonstrating an extrusion promoting function of NIPBL^27,28^. We surmise that the processivity effects are not simply the result of reduced cohesin loading in the absence of NIPBL. In an independent study, direct depletion of cohesin to the same degree as that observed upon NIBPL depletion had little effect on loop extrusion (Elphège Nora, personal communication). Therefore, our in vivo data are consistent with NIPBL’s ability to enhance cohesin extrusion, rather than merely serving as a cohesin loader onto chromatin.

Our findings demonstrate that NIPBL is dispensable for the formation of many CRE loops, regardless of their length. This was true even for CRE loops spanning similar distances as structural loops which were dependent on NIPBL. Thus, CRE loops can form independently of cohesin-mediated extrusion and structural loop formation. Consistent with this, NIPBL depletion had minimal effects on transcription reactivation during mitotic exit, with only a small subset of genes being affected. Therefore, the majority of regulatory interactions likely form via alternative mechanisms, such as transcription (co)factor oligomerization/clustering.

For example, we showed that that NIPBL is largely dispensable for most LDB1- and YY1-driven interactions. Furthermore, the loops mediated by these two architectural proteins do not overlap significantly, suggesting that LDB1 and YY1 operate through distinct extrusion-independent mechanisms^53^. However, LDB1- and YY1-dependent loops represent only a subset of all CRE interactions^52,53^, suggesting that additional (unidentified) factors are involved.

We found that the levels of H3K27ac at loop anchors were predicative of loop sensitivity to NIPBL depletion. In those cases in which loops with among weak CRE did form, they did so with delayed kinetics. Conversely, loops involving highly active enhancers exhibited similar formation kinetics in the presence and absence of NIPBL. This differential dependency upon NIPBL could result from lower transcription (co)factor concentrations at weak CREs, rendering them more reliant on loop extrusion to quickly facilitate long-range interactions.

In sum, by leveraging mitotic NIPBL degradation and synchronous release into G1, we directly tested the role of loop extrusion in the de novo establishment of chromatin architecture. While NIPBL is required for TAD formation, most regulatory interactions form independently. These findings reveal distinct extrusion-dependent and - independent mechanisms that organize chromatin architecture, uncoupling loop extrusion from affinity-based mechanisms in the formation of regulatory connectivity.

## EXTENDED DATA FIGURE LEGENDS

**Extended data Fig. 1:**
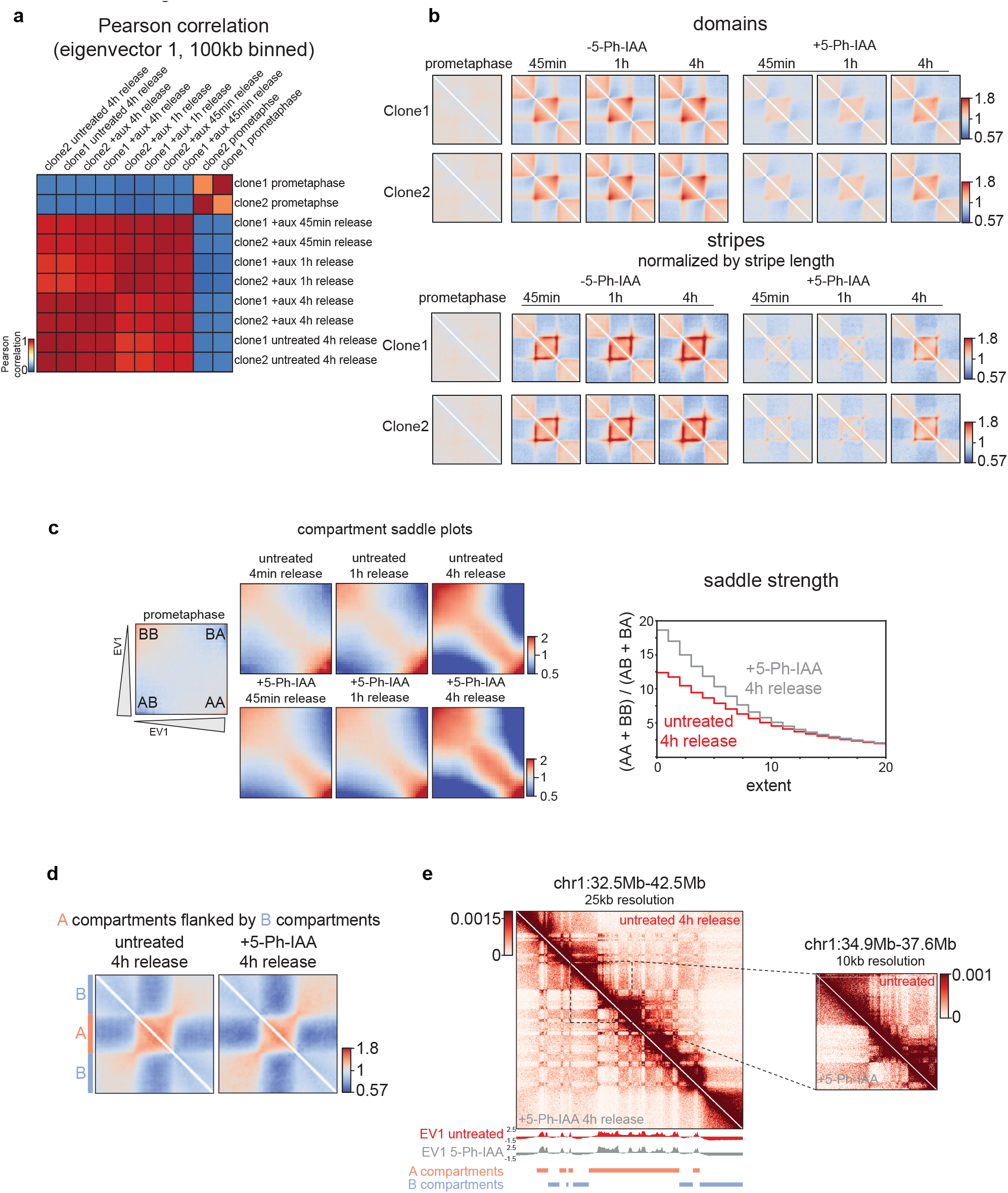
Hi-C replicate correlation and compartment analyses. **a**. Heatmap showing Pearson correlation coefficients for Hi-C samples based on eigenvector 1 values. **b**. (top) pile-up plots showing average TAD formation during G1 entry for each NIPBL-AID clone. (bottom) Pile-up plots showing average stripe formation (normalized by stripe length) for each NIPBL-AID clone. **c**. (left) saddle plots showing compartment strength during G1 entry at all timepoints. (right) quantification of compartment strength. **d**. Pile-up plots showing average contact frequencies for A compartments flanked by B compartments. **e**. Hi-C contact matrices showing representative region exhibiting increased compartmentalization upon NIPBL depletion.

**Extended data Fig. 2:**
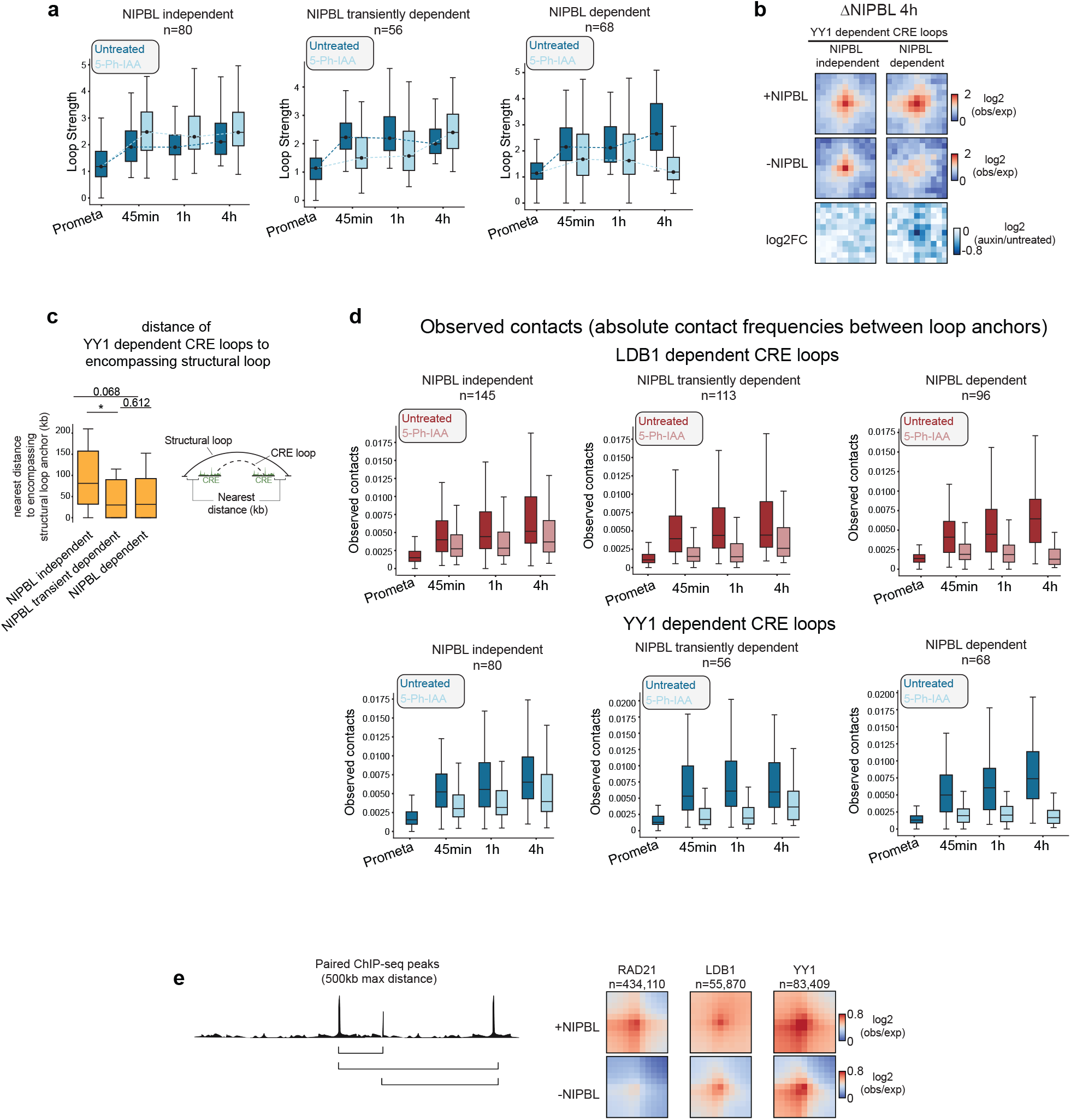
LDB1- and YY1-dependent loops can form in the absence of NIPBL. **a**. YY1-dependent CRE loop strength during G1 entry in the presence and absence of NIPBL. **b**. APA plots showing average signal for YY1-dependent CRE loops in the presence and absence of NIPBL (4h G1 condition). **c**. Distance of YY1-dependent CRE loops to encompassing structural loops. Only loops with identified structural loops encompassing them are shown. **d**. Observed contacts for LDB1- and YY1-dependent loops during G1 entry in the presence and absence of NIPBL. Loops stratified by NIPBL independent, delayed, and NIPBL dependent. These are the same loops from Extended Data Fig. 2a, and Fig. 4a. **e**. Paired chip-seq analysis. Asynchronous RAD21, LDB1, and YY1 peaks were paired to create all possible pairs of peaks within a maximum distance of 500kb. APA plots showing contact frequencies at late G1 (4h) in the presence and absence of NIPBL for each set of peak pairs.

## DATA AVAILABILITY

Hi-C and TT-seq datasets generated in this study are deposited in the GEO database (GSE256073). External data from prior studies listed here: LDB1, CTCF, RAD21, YY1, and H3K27ac ChIP-seq (GSE254377), LDB1 asynchronous Micro-C (GSE254373), YY1 asynchronous Micro-C (GSE247254).

## METHODS

### Nocodazole arrest/release and 5-Ph-IAA treatment

The G1E-ER4:Wapl^dTag^/Nipbl^mAID^ cells expressing OsTIR2 were treated with nocodazole (200ng/ml) for 8 hours at a density of 0.5 million/ml. To degrade Nipbl during mitotic arrest, 5-Ph-IAA (1uM) was added into the culture during the last 4 hours of nocodazole arrest. To acquire post mitotic samples without Nipbl, the cells were were released from nocodazole and resuspended in warm medium containing 5-Ph-IAA (1uM). The cells were allowed to enter G1 phase for 45 minutes, 1 hour or 4 hours respectively. Cells without 5-Ph-IAA treatment were collected in parallel as controls.

### Cell sorting

For *in-situ* Hi-C experiments, control and 5-Ph-IAA treated samples were acquired as described previously ^51^. Briefly, at the time of collection, cells were pelleted, washed and re-suspended in 1 x PBS. Cells were crosslinked with 2% formaldehyde for 10min at RT and subsequently quenched with 1M (final concentration) glycine for 5min at RT. Cells were pelleted and re-suspended in 1 x FACS sorting buffer (1 x PBS, 2% FBS, 2mM EDTA and 0.02% NaN3) containing 20ng/ml DAPI at a density of about 50-100 million cells/ml. G1 phase cells were sorted based on DAPI signal (2N). Sorted cells were snap-frozen and stored at −80°C.

### *In-situ* Hi-C

In-situ Hi-C was performed as previously described ^57^. Briefly, purified cells (0.1-0.2 million) were lysed in 1ml cold Cell Lysis Buffer (10mM Tris pH 8, 10mM NaCl, 0.2% NP-40) for 20min on ice. Subsequently, nuclei were washed pelleted and washed once with cold 1.4×NEB buffer 3.1 (NEB Cat#B7203S). Nuclei were then incubated in 25ul 1.4× NEB buffer 3.1 containing 0.1% SDS for 10min at 65°C. SDS was then quenched by 1% (final concentration) Triton X-100. Chromatin digest was performed by treatment with 25U DpnII restriction enzyme for overnight at 37°C. After overnight digestion, another stroke of DpnII (25U) was added into the system to further digest the genome for 4 hours at 37°C. DpnII heat inactivation was performed at 65°C for 20min. Digested chromatin was then repaired with 6.6U of DNA Polymerase I, Large (Klenow) fragment (NEB, Cat#M0210) in the presence of dCTP, dGTP, dTTP (Diamond Cat#B110049, B110050, B110051) and Biotin-14-dATP (active motif, Catalog No: 14138). Blunt end chromatin was then ligated with 2000 U of T4 DNA ligase (NEB, Cat#M0202M) for 4 hours at 16°C and 4 hours at room temperature. Chromatin was then reverse-crosslinked overnight at 65°C in the presence of 1% SDS and proteinase K (G-clone Cat#EZ0970). DNase-free RNase A (Vazyme Cat#DE111-01) was added to the system for 30min at 37°C to remove RNA. DNA was then extracted by phenol-chloroform extraction, precipitated, and dissolved in 130□l of nuclease free water. DNA was then sonicated to fragments of 200-300bp using a QSonica Q800R3 sonicator (40% amplitude, 15s ON and 15s OFF, 13min sonication). Fragmented DNA was purified through 1.8x VAHTS DNA Clean Beads (Vazyme Cat#N411). Biotin labeled DNA fragments were pulled down by Dynabeads MyOne Streptavidin C1 beads (Thermo Fisher Scientific, Cat#65002). DNA capturing streptavidin beads were washed twice with 1 X B&W buffer (5mM Tris-HCl, pH 7.5; 0.5mM EDTA; 1M NaCl) and subject to bead library preparation using VAHTS Universal DNA Library Prep Kit for MGI (Vazyme, Cat#NDM607-02) based on the manufacture’s protocol. Adaptor ligated DNA library was eluted in 0.1% SDS by incubating at 98°C for 10min. Finally, the library was amplified for 6-8 cycles on a thermal cycler, using the VAHTS® HiFi Amplification Mix. PCR products were then purified with 1.8x VAHTS DNA Clean Beads and sequenced on MGI DNBSEQ-T7 sequencing platform.

## Quantification and data analysis

### Loop calling and differential loop analysis

.hic files were converted to cool files using hic2cool (v0.8.3). Cooltools.dots was used on merged contact maps for each treatment condition to identify loops. We first identified loops using 10kb and 25kb resolutions. We used the following parameters to call dots: max_loci_separation=2_000_000, clustering_radius=20_000, lambda_bin_fdr=0.05, n_lambda_bins=50. We utilized rounded donut, vertical, horizontal and rounded lowleft kernels to define dots (pixels) that were enriched relative to local backgrounds.

We created a consensus loop list by: 1-merging loops identified in the untreated (4hr nocodazole release) and 5-Ph-IAA-treated (4hr nocodazole release) conditions for 10kb and 25kb resolutions, 2-merging untreated/5-Ph-IAA loops from each resolution, retaining the highest (smallest) resolution coordinates if a loop was called at both resolutions. We then calculated loop strength at all consensus loops for untreated and 5-Ph-IAA-treated Hi-C contact matrices by calculating the observed/locally-adjusted expected value. We calculated the locally-adjusted expected value by multiplying the expected value at the peak pixel by the sum of the observed contacts in the rounded donut region divided by the sum of the expected contacts in the rounded donut region. Loop strength was calculated for each loop using the resolution at which the loop was originally identified. Loops with NA, 0, or infinite strength values were removed to filter out those in sparse regions in either untreated/5-Ph-IAA-treated conditions. We then calculated a log2FC for each loop (5-Ph-IAA/untreated) and set a log2FC cutoff of −/+ 0.5 to define weakened/strengthened loops upon NIPBL depletion.

### Stratifying loops based on ChIP-seq peaks and CRE annotations

We stratified loops into different categories based on whether their anchors overlapped (by at least 1bp) CTCF/cohesin ChIP-seq peaks and/or CREs. We used a list of annotated CREs from our prior study^41^ based on H3K27ac ChIP-seq and distance to TSSs. Briefly, enhancers were defined as H3K27ac-enriched regions that were located at least 1kb away from a TSS. Promoters were defined as H3K27ac-enriched regions located within 1kb of a TSS. A list of asynchronous CTCF/cohesin co-occupied sites were used from another prior study^53^. Loops were defined as structural loops: CTCF/cohesin co-occupied sites in both anchors and CREs in 1 or 0 anchors, CRE loops: CREs in both anchors, CTCF/cohesin co-occupied sites in 1 or 0 anchors, and dual-function loops: CTCF/cohesin co-occupied sites in both anchors and CREs in both anchors.

### Quantifying asynchronous Micro-C loop strength

We used a list of asynchronous Micro-C loops from our prior study that had CREs in both anchors (this included dual-function and CRE categories)^53^. We calculated the dependence of each loop on LDB1^53^, YY1^52^ or NIPBL (this study) by utilizing contact maps (Micro-C for LDB1/YY1, Hi-C for NIPBL) generated in the presence and absence of each factor. We quantified loop strength for each loop as previously described using 10kb contact maps for each dataset. We filtered out loops that were in sparse regions in any dataset by removing NA, 0 or infinite strength values. We further filtered for loops with a minimum loop strength value of 1.2 in all datasets. Importantly, this conservative strategy resulted in fewer loops that were dependent on either LDB1 or YY1 than we previously reported. This is due to the fact that some LDB1/YY1-dependent loops may be in sparse regions in the NIPBL datasets or not present/weak in late G1. Additionally, short-range LDB1/YY1 dependent loops may be “missed” due to quantifying loop strength at 10kb resolution.

We defined LDB1/YY1-dependent loops as those exhibiting a loop strength log2 fold change (log2FC) of less than −0.5, corresponding to an approximate 30% reduction in loop strength following LDB1 or YY1 depletion. To asses their responses to NIPBL depletion, we then classified LDB/YY1 dependent loops into three categories based on loop strength comparisons between untreated and 5-Ph-IAA-treated conditions (−/+NIPBL) at 45min, 1hr and 4hr post-nocodazole release. NIPBL-independent loops-loops with log2FC values greater than −0.5 at all time points, indicating minimal influence of NIPBL. NIPBL-transiently affected loops-loops with log2FC values less than −0.5 at either the 45min or 1hr time points, but greater than −0.5 at the 4hr time point, suggesting a temporary reduction in loop strength due to NIPBL depletion. NIPBL-dependent loops-loops with log2FC values less than −0.5 at the 4hr timepoint, indicating that ultimately these loops could not form in the absence of NIPBL to the same degree as in the untreated condition.

### Identifying and Visualizing TADs and Stripes

We identified TADs using 10kb resolution contact maps using the HiCFindTADs function from HiCExplorer with default parameters. We generated Pileups of all TADs identified in the untreated 4hr nocodazole release condition using the python package Coolpup.

We identified stripes using 10kb resolution contact maps using Stripenn^47^, and generated pileups of all stripes from untreated 4hr nocodazole release with Coolpup.

### A/B compartment assignment and saddle plotting

We used cooltools (v0.7.1) to compute cis eigenvector values on 100kb binned Hi-C matrices from all NIPBL-AID treatment conditions. A compartments were identified as those with positive EV1 values, and B compartments as those with negative EV1 values. This assignment was based on GC content. We generated saddle plots using the cooltools.saddle function to show all AA, BB, AB, and BA interactions genome-wide.

### TT-seq differential gene analysis

Strand-spec TT-seq reads were quantified in gene bodies using deepTools v3.5.5. We quantified raw, mapped reads in positively-stranded genes using reads from the forward strand and vice versa for negatively-stranded genes. We used the longest transcript of Refseq annotated, mm9 genes. We then used DESEQ2 to perform differential expression analysis.

## Acknowledgements

This work was supported by grants 1F31DK136200-01A1, T32GM008216 and the Blavatnik Family Fellowship Award to N.G.A.; National Science Foundation of China Grant 321004422 to H.Z.; and R01DK05937, R01DK058044, and U01DK127405 to G.A.B.

## References

1. Vermunt, M.W., Zhang, D., and Blobel, G.A. (2019). The interdependence of gene-regulatory elements and the 3D genome. J Cell Biol 218, 12–26. 10.1083/jcb.201809040.

2. Panigrahi, A., and O’Malley, B.W. (2021). Mechanisms of enhancer action: the known and the unknown. Genome Biol 22, 108. 10.1186/s13059-021-02322-1.

3. Rao, S.S., Huntley, M.H., Durand, N.C., Stamenova, E.K., Bochkov, I.D., Robinson, J.T., Sanborn, A.L., Machol, I., Omer, A.D., Lander, E.S., and Aiden, E.L. (2014). A 3D map of the human genome at kilobase resolution reveals principles of chromatin looping. Cell 159, 1665–1680. 10.1016/j.cell.2014.11.021.

4. Hansen, A.S. (2020). CTCF as a boundary factor for cohesin-mediated loop extrusion: evidence for a multi-step mechanism. Nucleus 11, 132–148. 10.1080/19491034.2020.1782024.

5. de Wit, E., and Nora, E.P. (2023). New insights into genome folding by loop extrusion from inducible degron technologies. Nat Rev Genet 24, 73–85. 10.1038/s41576-022-00530-4.

6. Fudenberg, G., Imakaev, M., Lu, C., Goloborodko, A., Abdennur, N., and Mirny, L.A. (2016). Formation of Chromosomal Domains by Loop Extrusion. Cell Rep 15, 2038–2049. 10.1016/j.celrep.2016.04.085.

7. Dixon, J.R., Selvaraj, S., Yue, F., Kim, A., Li, Y., Shen, Y., Hu, M., Liu, J.S., and Ren, B. (2012). Topological domains in mammalian genomes identified by analysis of chromatin interactions. Nature 485, 376–380. 10.1038/nature11082.

8. Nora, E.P., Lajoie, B.R., Schulz, E.G., Giorgetti, L., Okamoto, I., Servant, N., Piolot, T., van Berkum, N.L., Meisig, J., Sedat, J., et al. (2012). Spatial partitioning of the regulatory landscape of the X-inactivation centre. Nature 485, 381–385. 10.1038/nature11049.

9. Hou, C., Li, L., Qin, Z.S., and Corces, V.G. (2012). Gene density, transcription, and insulators contribute to the partition of the Drosophila genome into physical domains. Mol Cell 48, 471–484. 10.1016/j.molcel.2012.08.031.

10. Sexton, T., Yaffe, E., Kenigsberg, E., Bantignies, F., Leblanc, B., Hoichman, M., Parrinello, H., Tanay, A., and Cavalli, G. (2012). Three-dimensional folding and functional organization principles of the Drosophila genome. Cell 148, 458–472. 10.1016/j.cell.2012.01.010.

11. Beagan, J.A., and Phillips-Cremins, J.E. (2020). On the existence and functionality of topologically associating domains. Nat Genet 52, 8–16. 10.1038/s41588-019-0561-1.

12. Gabriele, M., Brandao, H.B., Grosse-Holz, S., Jha, A., Dailey, G.M., Cattoglio, C., Hsieh, T.S., Mirny, L., Zechner, C., and Hansen, A.S. (2022). Dynamics of CTCF- and cohesin-mediated chromatin looping revealed by live-cell imaging. Science 376, 496–501. 10.1126/science.abn6583.

13. Sabaté, T., Lelandais, B., Robert, M.-C., Szalay, M., Tinevez, J.-Y., Bertrand, E., and Zimmer, C. (2024). Universal dynamics of cohesin-mediated loop extrusion. bioRxiv.

14. Haarhuis, J.H.I., van der Weide, R.H., Blomen, V.A., Yanez-Cuna, J.O., Amendola, M., van Ruiten, M.S., Krijger, P.H.L., Teunissen, H., Medema, R.H., van Steensel, B., et al. (2017). The Cohesin Release Factor WAPL Restricts Chromatin Loop Extension. Cell 169, 693–707 e614. 10.1016/j.cell.2017.04.013.

15. Davidson, I.F., and Peters, J.M. (2021). Genome folding through loop extrusion by SMC complexes. Nat Rev Mol Cell Biol 22, 445–464. 10.1038/s41580-021-00349-7.

16. Tedeschi, A., Wutz, G., Huet, S., Jaritz, M., Wuensche, A., Schirghuber, E., Davidson, I.F., Tang, W., Cisneros, D.A., Bhaskara, V., et al. (2013). Wapl is an essential regulator of chromatin structure and chromosome segregation. Nature 501, 564–568. 10.1038/nature12471.

17. Ouyang, Z., Zheng, G., Tomchick, D.R., Luo, X., and Yu, H. (2016). Structural Basis and IP6 Requirement for Pds5-Dependent Cohesin Dynamics. Mol Cell 62, 248–259. 10.1016/j.molcel.2016.02.033.

18. Wutz, G., Varnai, C., Nagasaka, K., Cisneros, D.A., Stocsits, R.R., Tang, W., Schoenfelder, S., Jessberger, G., Muhar, M., Hossain, M.J., et al. (2017). Topologically associating domains and chromatin loops depend on cohesin and are regulated by CTCF, WAPL, and PDS5 proteins. EMBO J 36, 3573–3599. 10.15252/embj.201798004.

19. Rollins, R.A., Korom, M., Aulner, N., Martens, A., and Dorsett, D. (2004). Drosophila nipped-B protein supports sister chromatid cohesion and opposes the stromalin/Scc3 cohesion factor to facilitate long-range activation of the cut gene. Mol Cell Biol 24, 3100–3111. 10.1128/MCB.24.8.3100-3111.2004.

20. Schwarzer, W., Abdennur, N., Goloborodko, A., Pekowska, A., Fudenberg, G., Loe-Mie, Y., Fonseca, N.A., Huber, W., Haering, C.H., Mirny, L., and Spitz, F. (2017). Two independent modes of chromatin organization revealed by cohesin removal. Nature 551, 51–56. 10.1038/nature24281.

21. Gillespie, P.J., and Hirano, T. (2004). Scc2 couples replication licensing to sister chromatid cohesion in Xenopus egg extracts. Curr Biol 14, 1598–1603. 10.1016/j.cub.2004.07.053.

22. Watrin, E., Schleiffer, A., Tanaka, K., Eisenhaber, F., Nasmyth, K., and Peters, J.M. (2006). Human Scc4 is required for cohesin binding to chromatin, sister-chromatid cohesion, and mitotic progression. Curr Biol 16, 863–874. 10.1016/j.cub.2006.03.049.

23. Murayama, Y., and Uhlmann, F. (2014). Biochemical reconstitution of topological DNA binding by the cohesin ring. Nature 505, 367–371. 10.1038/nature12867.

24. Kurokawa, Y., and Murayama, Y. (2020). DNA Binding by the Mis4(Scc2) Loader Promotes Topological DNA Entrapment by the Cohesin Ring. Cell Rep 33, 108357. 10.1016/j.celrep.2020.108357.

25. Gutierrez-Escribano, P., Newton, M.D., Llauro, A., Huber, J., Tanasie, L., Davy, J., Aly, I., Aramayo, R., Montoya, A., Kramer, H., et al. (2019). A conserved ATP-and Scc2/4-dependent activity for cohesin in tethering DNA molecules. Sci Adv 5, eaay6804. 10.1126/sciadv.aay6804.

26. Alonso-Gil, D., and Losada, A. (2023). NIPBL and cohesin: new take on a classic tale. Trends Cell Biol 33, 860–871. 10.1016/j.tcb.2023.03.006.

27. Davidson, I.F., Bauer, B., Goetz, D., Tang, W., Wutz, G., and Peters, J.M. (2019). DNA loop extrusion by human cohesin. Science 366, 1338–1345. 10.1126/science.aaz3418.

28. Kim, Y., Shi, Z., Zhang, H., Finkelstein, I.J., and Yu, H. (2019). Human cohesin compacts DNA by loop extrusion. Science 366, 1345–1349. 10.1126/science.aaz4475.

29. Alonso-Gil, D., Cuadrado, A., Gimenez-Llorente, D., Rodriguez-Corsino, M., and Losada, A. (2023). Different NIPBL requirements of cohesin-STAG1 and cohesin-STAG2. Nat Commun 14, 1326. 10.1038/s41467-023-36900-7.

30. Petela, N.J., Gligoris, T.G., Metson, J., Lee, B.G., Voulgaris, M., Hu, B., Kikuchi, S., Chapard, C., Chen, W., Rajendra, E., et al. (2018). Scc2 Is a Potent Activator of Cohesin’s ATPase that Promotes Loading by Binding Scc1 without Pds5. Mol Cell 70, 1134–1148 e1137. 10.1016/j.molcel.2018.05.022.

31. Thiecke, M.J., Wutz, G., Muhar, M., Tang, W., Bevan, S., Malysheva, V., Stocsits, R., Neumann, T., Zuber, J., Fraser, P., et al. (2020). Cohesin-Dependent and - Independent Mechanisms Mediate Chromosomal Contacts between Promoters and Enhancers. Cell Rep 32, 107929. 10.1016/j.celrep.2020.107929.

32. Rao, S.S.P., Huang, S.C., Glenn St Hilaire, B., Engreitz, J.M., Perez, E.M., Kieffer-Kwon, K.R., Sanborn, A.L., Johnstone, S.E., Bascom, G.D., Bochkov, I.D., et al. (2017). Cohesin Loss Eliminates All Loop Domains. Cell 171, 305–320 e324. 10.1016/j.cell.2017.09.026.

33. Nora, E.P., Goloborodko, A., Valton, A.L., Gibcus, J.H., Uebersohn, A., Abdennur, N., Dekker, J., Mirny, L.A., and Bruneau, B.G. (2017). Targeted Degradation of CTCF Decouples Local Insulation of Chromosome Domains from Genomic Compartmentalization. Cell 169, 930–944 e922. 10.1016/j.cell.2017.05.004.

34. Hsieh, T.S., Cattoglio, C., Slobodyanyuk, E., Hansen, A.S., Darzacq, X., and Tjian, R. (2022). Enhancer-promoter interactions and transcription are largely maintained upon acute loss of CTCF, cohesin, WAPL or YY1. Nat Genet 54, 1919–1932. 10.1038/s41588-022-01223-8.

35. Rhodes, J.D.P., Feldmann, A., Hernandez-Rodriguez, B., Diaz, N., Brown, J.M., Fursova, N.A., Blackledge, N.P., Prathapan, P., Dobrinic, P., Huseyin, M.K., et al. (2020). Cohesin Disrupts Polycomb-Dependent Chromosome Interactions in Embryonic Stem Cells. Cell Rep 30, 820–835 e810. 10.1016/j.celrep.2019.12.057.

36. Vian, L., Pekowska, A., Rao, S.S.P., Kieffer-Kwon, K.R., Jung, S., Baranello, L., Huang, S.C., El Khattabi, L., Dose, M., Pruett, N., et al. (2018). The Energetics and Physiological Impact of Cohesin Extrusion. Cell 175, 292–294. 10.1016/j.cell.2018.09.002.

37. Naumova, N., Imakaev, M., Fudenberg, G., Zhan, Y., Lajoie, B.R., Mirny, L.A., and Dekker, J. (2013). Organization of the mitotic chromosome. Science 342, 948–953. 10.1126/science.1236083.

38. Gibcus, J.H., Samejima, K., Goloborodko, A., Samejima, I., Naumova, N., Nuebler, J., Kanemaki, M.T., Xie, L., Paulson, J.R., Earnshaw, W.C., et al. (2018). A pathway for mitotic chromosome formation. Science 359. 10.1126/science.aao6135.

39. Ito, K., and Zaret, K.S. (2022). Maintaining Transcriptional Specificity Through Mitosis. Annu Rev Genomics Hum Genet 23, 53–71. 10.1146/annurev-genom-121321-094603.

40. Kadauke, S., and Blobel, G.A. (2013). Mitotic bookmarking by transcription factors. Epigenetics Chromatin 6, 6. 10.1186/1756-8935-6-6.

41. Zhang, H., Emerson, D.J., Gilgenast, T.G., Titus, K.R., Lan, Y., Huang, P., Zhang, D., Wang, H., Keller, C.A., Giardine, B., et al. (2019). Chromatin structure dynamics during the mitosis-to-G1 phase transition. Nature 576, 158–162. 10.1038/s41586-019-1778-y.

42. Nishimura, K., Fukagawa, T., Takisawa, H., Kakimoto, T., and Kanemaki, M. (2009). An 5-Ph-IAA-based degron system for the rapid depletion of proteins in nonplant cells. Nat Methods 6, 917–922. 10.1038/nmeth.1401.

43. Natsume, T., Kiyomitsu, T., Saga, Y., and Kanemaki, M.T. (2016). Rapid Protein Depletion in Human Cells by 5-Ph-IAA-Inducible Degron Tagging with Short Homology Donors. Cell Rep 15, 210–218. 10.1016/j.celrep.2016.03.001.

44. Weiss, M.J., Yu, C., and Orkin, S.H. (1997). Erythroid-cell-specific properties of transcription factor GATA-1 revealed by phenotypic rescue of a gene-targeted cell line. Mol Cell Biol 17, 1642–1651. 10.1128/MCB.17.3.1642.

45. Ramirez, F., Bhardwaj, V., Arrigoni, L., Lam, K.C., Gruning, B.A., Villaveces, J., Habermann, B., Akhtar, A., and Manke, T. (2018). High-resolution TADs reveal DNA sequences underlying genome organization in flies. Nat Commun 9, 189. 10.1038/s41467-017-02525-w.

46. Zhao, H., Lin, Y., Lin, E., Liu, F., Shu, L., Jing, D., Wang, B., Wang, M., Shan, F., Zhang, L., et al. (2024). Genome folding principles uncovered in condensin-depleted mitotic chromosomes. Nat Genet 56, 1213–1224. 10.1038/s41588-024-01759-x.

47. Yoon, S., Chandra, A., and Vahedi, G. (2022). Stripenn detects architectural stripes from chromatin conformation data using computer vision. Nat Commun 13, 1602. 10.1038/s41467-022-29258-9.

48. Barrington, C., Georgopoulou, D., Pezic, D., Varsally, W., Herrero, J., and Hadjur, S. (2019). Enhancer accessibility and CTCF occupancy underlie asymmetric TAD architecture and cell type specific genome topology. Nat Commun 10, 2908. 10.1038/s41467-019-10725-9.

49. Liu, Y., and Dekker, J. (2022). CTCF-CTCF loops and intra-TAD interactions show differential dependence on cohesin ring integrity. Nat Cell Biol 24, 1516–1527. 10.1038/s41556-022-00992-y.

50. Nuebler, J., Fudenberg, G., Imakaev, M., Abdennur, N., and Mirny, L.A. (2018). Chromatin organization by an interplay of loop extrusion and compartmental segregation. Proc Natl Acad Sci U S A 115, E6697–E6706. 10.1073/pnas.1717730115.

51. Zhang, H., Lam, J., Zhang, D., Lan, Y., Vermunt, M.W., Keller, C.A., Giardine, B., Hardison, R.C., and Blobel, G.A. (2021). CTCF and transcription influence chromatin structure re-configuration after mitosis. Nat Commun 12, 5157. 10.1038/s41467-021-25418-5.

52. Lam, J.C., Aboreden, N.G., Midla, S.C., Wang, S., Huang, A., Keller, C.A., Giardine, B., Henderson, K.A., Hardison, R.C., Zhang, H., and Blobel, G.A. (2024). YY1-controlled regulatory connectivity and transcription are influenced by the cell cycle. Nat Genet 56, 1938–1952. 10.1038/s41588-024-01871-y.

53. Aboreden, N.G., Lam, J.C., Goel, V.Y., Wang, S., Wang, X., Midla, S.C., Quijano, A., Keller, C.A., Giardine, B.M., Hardison, R.C., et al. (2024). LDB1 establishes multi-enhancer networks to regulate gene expression. Mol Cell. 10.1016/j.molcel.2024.11.037.

54. Cuartero, S., Weiss, F.D., Dharmalingam, G., Guo, Y., Ing-Simmons, E., Masella, S., Robles-Rebollo, I., Xiao, X., Wang, Y.F., Barozzi, I., et al. (2018). Control of inducible gene expression links cohesin to hematopoietic progenitor self-renewal and differentiation. Nat Immunol 19, 932–941. 10.1038/s41590-018-0184-1.

55. Hsiung, C.C., Bartman, C.R., Huang, P., Ginart, P., Stonestrom, A.J., Keller, C.A., Face, C., Jahn, K.S., Evans, P., Sankaranarayanan, L., et al. (2016). A hyperactive transcriptional state marks genome reactivation at the mitosis-G1 transition. Genes Dev 30, 1423–1439. 10.1101/gad.280859.116.

56. Krivega, I., and Dean, A. (2017). LDB1-mediated enhancer looping can be established independent of mediator and cohesin. Nucleic Acids Res 45, 8255–8268. 10.1093/nar/gkx433.

57. Zhao, H., Lin, Y., Lin, E., Liu, F., Shu, L., Jing, D., Wang, B., Wang, M., Shan, F., Zhang, L., et al. (2024). Genome folding principles uncovered in condensin-depleted mitotic chromosomes. Nat Genet. 10.1038/s41588-024-01759-x.

